# Histone Tail Dynamics in Partially Disassembled Nucleosomes During Chromatin Remodeling

**DOI:** 10.1101/633370

**Authors:** T. Kameda, A. Awazu, Y. Togashi

**Affiliations:** Hiroshima University

## Abstract

Nucleosomes are structural units of the chromosome consisting of DNA wrapped around histone proteins, and play important roles in compaction and regulation of the chromatin structure. While the structure and dynamics of canonical nucleosomes have been studied extensively, those of nucleosomes in intermediate states, that occur when their structure or positioning is modulated, have been less understood. In particular, the dynamic features of partially disassembled nucleosomes have not been discussed in previous studies. Using all-atom molecular dynamics simulations, in this study, we investigated the dynamics and stability of nucleosome structures lacking a histone-dimer. DNA in nucleosomes lacking a histone H2A/H2B dimer was drastically deformed due to loss of local interactions between DNA and histones. In contrast, conformation of DNA in nucleosomes lacking H3/H4 was similar to the canonical nucleosome, as the H2A C-terminal domain infiltrated the space originally occupied by the dissociated H3/H4 histones and stabilized DNA in close proximity. Our results suggest that, besides histone chaperones, the intrinsic dynamics of nucleosomes support the exchange of H2A/H2B, which is significantly more frequent than that of H3/H4.

**Statement of Significance:** Eukaryotic chromosomes are composed of nucleosomes, in which the DNA wraps around the core histone proteins. To enable transcription and replication of DNA, or to modulate these functions by exchange of histones, nucleosomes should be at least partially disassembled, as evidenced by the observation of nucleosome structures lacking an H2A/H2B histone dimer by crystallography. The dynamic behavior of nucleosomes in such intermediate states may affect gene expression and repair, however it has not been completely elucidated so far. In this study, we adopted molecular dynamics simulations to analyze the conformational changes in partially disassembled nucleosomes. Enhanced structural fluctuations of DNA were observed in these nucleosomes, which may, as well as specific histone chaperones, support the exchange of H2A/H2B.

## Introduction

Eukaryotic genomic DNA is compacted into a hierarchical chromatin structure that is confined within the nucleus. Basic units constituting chromatin are complexes of DNA and histone proteins, called nucleosomes (1). A nucleosome consists of approximately 147 base pairs (bps) of DNA and 8 histone proteins, namely H2A, H2B, H3, and H4. In the canonical nucleosome structure, two H2A/H2B and two H3/H4 hetero-dimers form a histone octamer (1). The DNA wraps around the histone proteins in the nucleosome, forming a spool-like structure. Amino-acid sequences of these histones are highly conserved among eukaryotes. The nucleosomal structures are stable and are supported by electrostatic interactions between DNA and histone.

Nucleosomes are strongly associated with chromatin structure formation and the genome function. Gene expression involves nucleosomal rearrangement. In turn, changes in nucleosome positioning regulate the accessibility of proteins to DNA and hence modulate gene expression, which is also regulated based largely on cell species and cell cycle (2–4). Eviction of histones and reconstruction of nucleosomes occur frequently upon rearrangement. These structural changes have been associated with the interaction of nucleosome and histone chaperones (5–7), and often occur upon transcription. FACT (FAcilitates Chromatin Transcription) is a well-known histone chaperone and is highly conserved among eukaryotes (7–9). Recent reports have postulated mechanisms of histone chaperones interacting with nucleosomes (10–12).

In these processes, nucleosomes may transiently lack some histone proteins (e.g. hexasome (13)). For example, during transcription, nucleosomes must at least be repositioned (14). Consequently, structural conflicts between neighboring nucleosomes may result in partial histone dissociation, manifesting *in vitro* as overlapping dinucleosomes (15). These partially disassembled nucleosomes (intermediate nucleosomes; or “partially assembled nucleosome” coined by Rychkov *et al*. (16)) have been reported in nucleosome structure modulation, histone assembly and disassembly (6, 7, 10, 13). For example, the histone variant H2A.Z is widely distributed in higher eukaryotic genomes and regulates genome dynamics (17, 18). H2A.Z is inserted into H2A/H2B-lacking nucleosomes with the help of histone chaperones. In some cases, partially disassembled nucleosomes facilitate binding of proteins to DNA. Such attachment and detachment of histones occur over the entire chromosomes, and partially disassembled nucleosomes may occur upon detachment of any of the constituent histones (19). In fact, the dissociation frequency of histone H2A/H2B is significantly higher than that of H3/H4 (20), and nucleosomes lacking only H3/H4 have not been experimentally observed. Therefore, most intermediate nucleosomes are considered to be hexasomes lacking H2A/H2B (13).

Considering their importance, the structures of these intermediate nucleosomes have been extensively explored of late by crystallography and cryo-EM methods. However, the structural dynamics, with respect to histone dissociation, have not been fully elucidated. In this research, we focused on the intermediate nucleosome dynamics using molecular dynamics (MD) simulation. We considered nucleosome structures that lack one histone dimer, as well as the canonical nucleosome, and analyzed the differences in their dynamics. Deletion of a histone dimer had only little influence on other histones. In contrast, DNA dynamics was drastically affected, based on the eliminated histone type. These results are consistent with previous findings in nucleosome modulation, and suggest the importance of transient dynamics of intermediate nucleosomes in chromatin remodeling.

## Materials and Methods

### Nucleosome Models and Simulation Procedures

We employed all-atom MD simulations for nucleosome dynamics in solution. We constructed 5 nucleosome models, using the crystal structure of a nucleosome (PDB ID: 1KX5) (1). The canonical nucleosome corresponds to the entire crystal structure. The others are models of partially disassembled nucleosomes, constructed by removing one of the two H2A/H2B hetero-dimers or the two H3/H4 hetero-dimers from the canonical nucleosome; which we named **ΔH3/H4, ΔH2A/H2B, ΔH3**′/**H4**′, and **ΔH2A**′/**H2B**′ (Fig. 1; these histones are indexed as chains A to H in the PDB file). Then, these models were soaked into a 153 Å × 197 Å × 103 Å water box with 150 mM KCl. TIP3P water model was employed. All histidine residues were configured as *ϵ*-protonated. VMD (21) was used to infer missing atom coordinates, solvate the model, and visualize the structure throughout the study.

**Figure 1:**
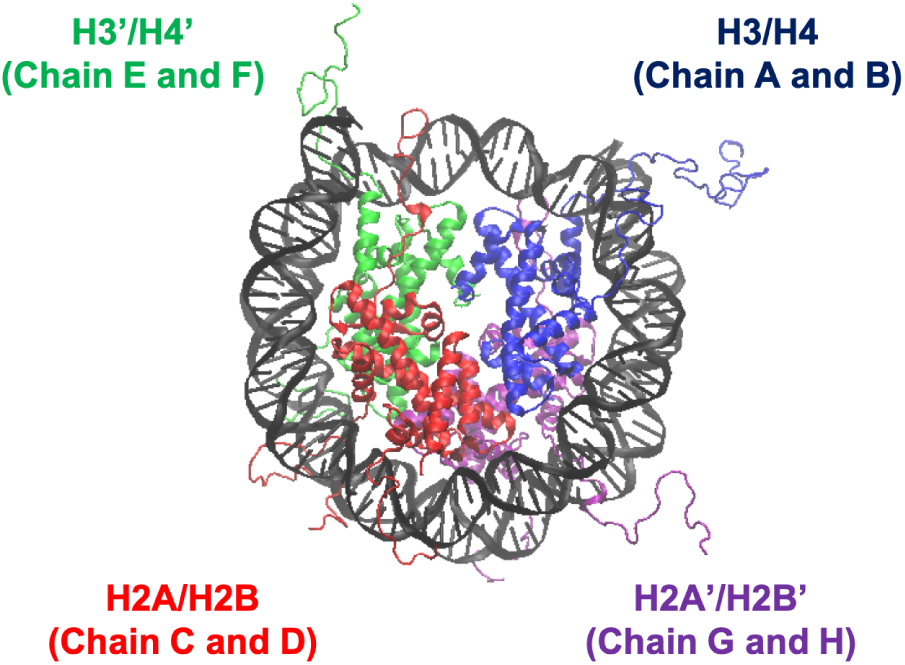
Nucleosome Structure and Histone Indices. PDB ID: 1KX5. Histones H3/H4 (Chain A and B), H2A/H2B (Chain C and D), H3′/H4′ (Chain E and F), and H2A′/H2B′ (Chain G and H) are shown in blue, red, green, and purple, respectively. Following these indices, we named each nucleosome lacking one of the histone-dimers as Δ H3/H4, Δ H2A/H2B, Δ H3’/H4’, and Δ H2A’/H2B’.

All simulations were carried out using NAMD (version 2.12 multi-core with CUDA) (22). CHARMM36 force-field was used. Periodic boundary condition with Particle-Mesh Ewald electrostatics was employed. Cutoff 12 Å (with switching from 10 Å) was used for non-bonded interactions. Temperature and pressure were set at 310 K and 1 atm, respectively; Langevin thermostat (damping coefficient: 5/ps) and Langevin-piston barostat were adopted. After energy minimization with harmonic restraints on C^*α*^ of amino acids and C1’ of nucleotides (spring constant: 100 pN/Å; 10000 steps), the system was simulated in equilibrium for 100 ns (time-step: 2 fs). The structures were sampled at 10 ps intervals, and each nucleosome model was simulated 10 times.

### Root Mean Square Displacement (RMSD)

The coordinate of atom *i* is denoted by ***x***_*i*_(*t*). We used heavy atoms except the histone tails (Table 1) for calculation, as these tails are highly disordered (same for RMSF and PCA). First, we aligned the nucleosome by eliminating the translational and rotational motions to minimize RMSD (i), and then calculated RMSD. Additionally, to evaluate the structural changes of DNA, we aligned the nucleosome according to only the core regions of histones (i.e. except DNA) (ii), and calculated RMSD likewise. Below, the coordinates after the alignment in the case of (i) and (ii) are denoted by 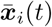 and 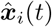, respectively.

**Table 1:** Residue IDs of Core Region and Tail Region for each Histones. Each chain ID and residue ID refer to PDB ID: 1KX5 (1). Only histone H2A has a C-terminal tail.

| Histone (Chain ID) | Core Region | Tail Region |
| --- | --- | --- |
| H3 (A) and H3' (E) | 45 – 135 | 1 – 44 |
| H4 (B) and H4' (F) | 25 – 102 | 1 – 24 |
| H2A (C) and H2A' (G) | 18 – 98 | 1 – 17 (N-terminal)<br>99 – 128 (C-terminal) |
| H2B (D) and H2B' (H) | 35 – 122 | 1 – 34 |

### Root Mean Square Fluctuation (RMSF) of DNA

Root mean square fluctuation of nucleotide *i* was defined as 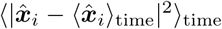. In this case, 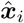 is the C1’ coordinate of nucleotide *i* after alignment. Note that ⟨·⟩_time_ denotes averaging over the latter half of the simulation (50.01 – 100 ns) and 10 trials for the same nucleosome model (same for below).

### Principal Component Analysis (PCA) of DNA

Principal component analysis was employed to analyze histone-wrapped DNA dynamics. PCA is often used in similar cases to evaluate protein dynamics (23). In the current case, each nucleotide was represented by the C1’ coordinate. Displacement *S*(*τ*, mode) := (*X*(*τ*) − ⟨*X*⟩_time_) · **v**_mode_ was evaluated where 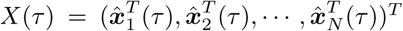, and **v**_*j*_ is *j*-th eigenvector of covariance matrix of *X*.

### Secondary Structure Determination

To determine the secondary structures in the histones, we employed the DSSP program (24). In the canonical nucleosome, each histone is composed of only several *α*-helices (1). We determined *α*-helix formation and the occurrence at each residue was calculated for the latter half of the simulation trajectory.

### Contact Frequency of Nucleotides and Amino Acids

Contact frequency between DNA and histone tails were evaluated as follows. The distance between C1’ of each nucleotide and C^*α*^ of each amino acid was examined. The molecules were regarded in contact if the distance was less than 10 Å. The contact frequency, defined as the ratio of samples in which the pair was in contact, was averaged over the latter half of the simulation and 10 trials for each nucleosome model.

## Results

### Structures of Histone Proteins

First, we analyzed the effects of partial histone dissociation on the secondary structure of other histones. The occurrence ratio of *α*-helix formation is shown in Fig. 2 and Fig. S1 in the Supporting Material. Each nucleosome lacking one histone dimer shows a profile of secondary structure formation similar to the canonical nucleosome. Removal of a histone dimer has little influence on the secondary structure of other remaining histone proteins. On the other hand, *α*-helix formation was occasionally observed in histone tail residues; which agrees with previous reports suggesting that histone-tail regions show transient *α*-helix formation when interacting with DNA (25, 26).

**Figure 2:**
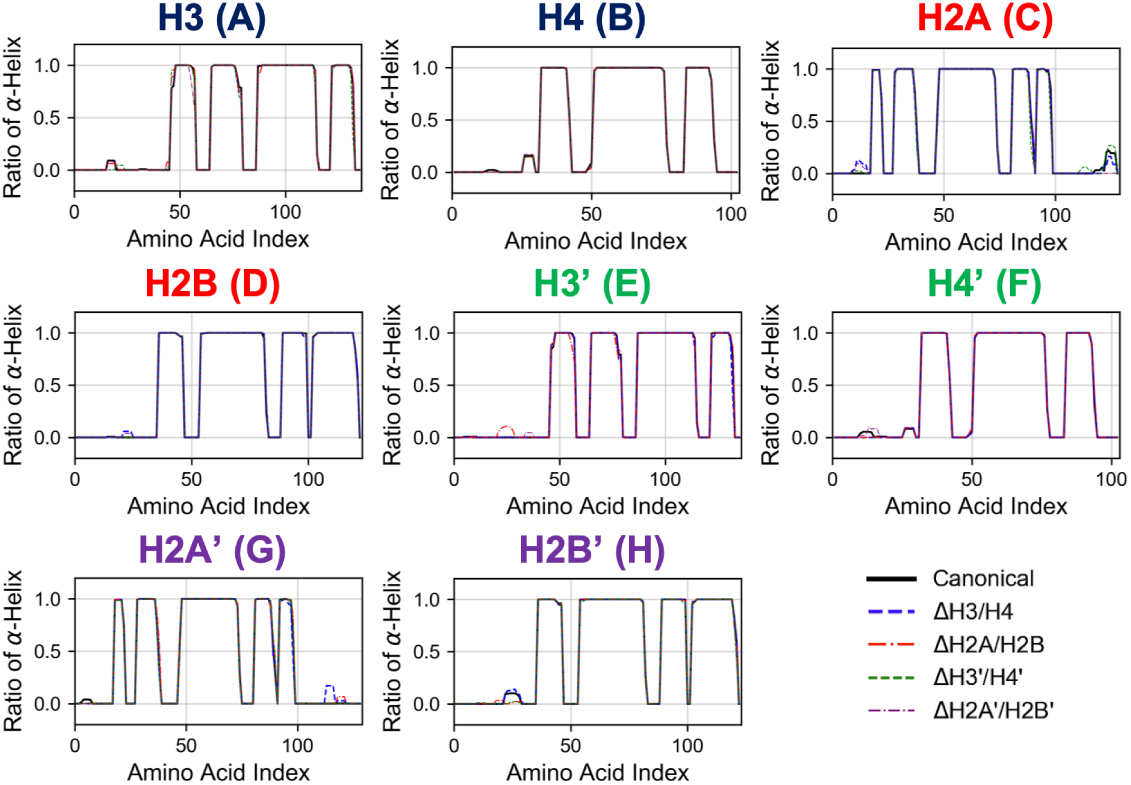
Secondary Structure Stability of Histones in Each Nucleosome. Averaged over 10 trials (each trial is shown in Fig. S1). Horizontal and vertical axes of each graph show amino acid index and ratio of *α*-helix formation in respective histones as indicated in figure.

Next, we evaluated the time series of RMSD of histones in each nucleosome, to analyze their tertiary structures (Figs. 3 and S2). In this case, RMSD was defined as 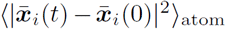, where ⟨⟩_atom_ means averaging over the constituent heavy atoms. Similar to the results of secondary structure analysis, each histone displayed similar profiles across all nucleosome models. Again, the dissociation of a histone dimer induced negligible effect on the other histones, which is consistent also with experimental observations (13, 27).

**Figure 3:**
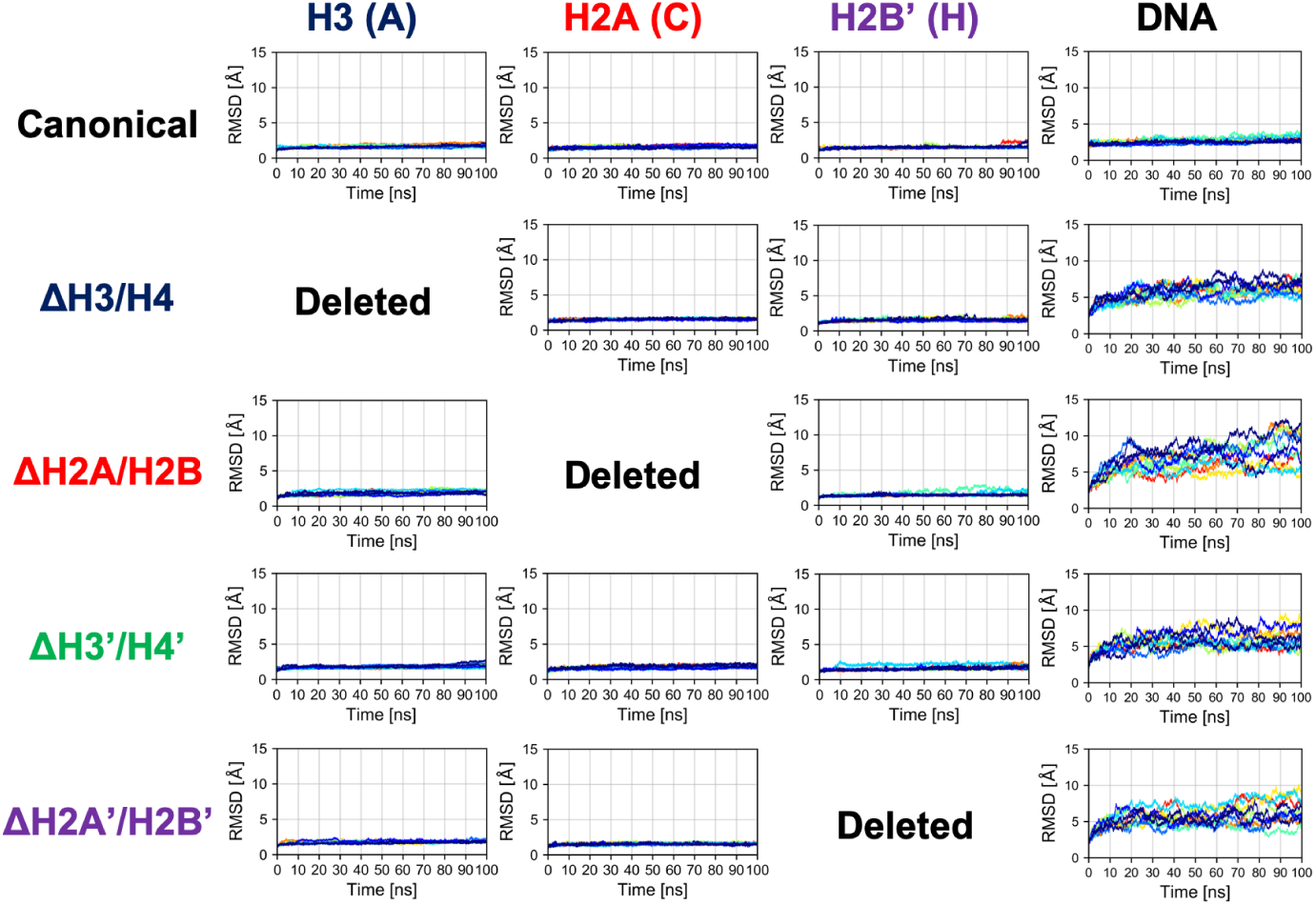
Time Series of RMSD of Histone and DNA in Each Nucleosome. Colors show different simulation trajectories (total 10).

### Structural Deformation of Nucleosomal DNA

In contrast, as shown in Fig. 3, histone-wrapped DNA dynamics was dramatically affected. RMSD of DNA in nucleosomes lacking any histone dimer was increased, compared to the canonical nucleosome; i.e., DNA was deformed from the native structure by the partial disassembly.

Then, we evaluated the deformation in each part of DNA, to determine which part was affected by the loss of histones. RMSF profiles of DNA are shown in Fig. S3. Fluctuations were increased only in nucleotides adjacent to the removed histone. Consistently, drastic deformations (superior modes of PCA) were observed in the region that lost interactions with histones (Fig. 4). These deformations mostly correspond to the breathing motion of DNA outward from the nucleosome.

**Figure 4:**
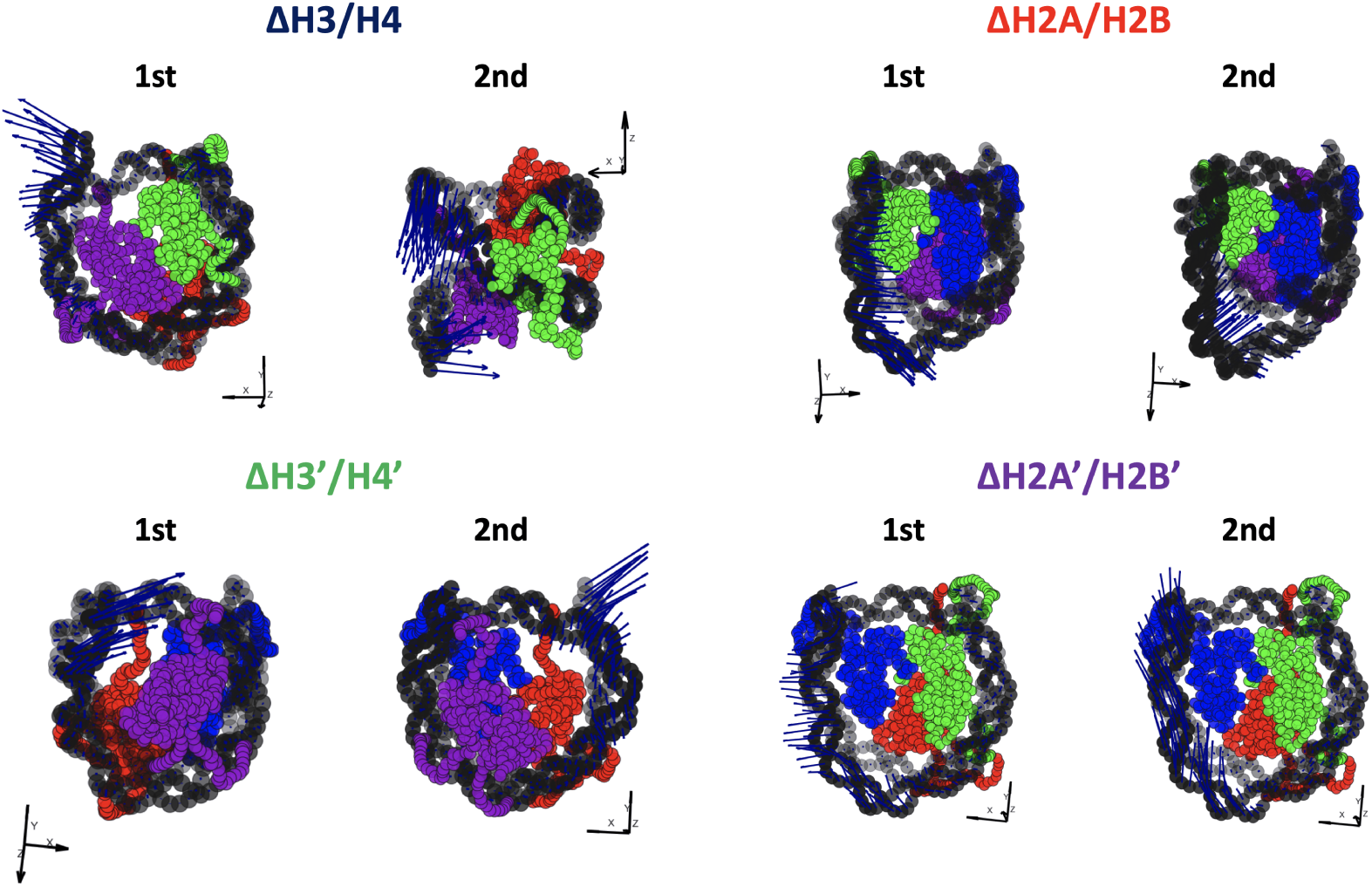
Effective Directions of DNA Deformation. Nucleotide (C1’ of each nucleotide) and amino acid (C^*α*^ of each amino acid) coordinates are shown by spheres. Blue arrows show the vectors of the 1st and 2nd PC modes. Arrow lengths indicate the norm of the vector.

To capture effective structural changes of DNA, we estimated free energy profiles projected to principal modes obtained from PCA analysis (see Materials and Methods). We also focused on trajectories projected onto this space, to evaluate DNA dynamics (Fig. 5).

**Figure 5:**
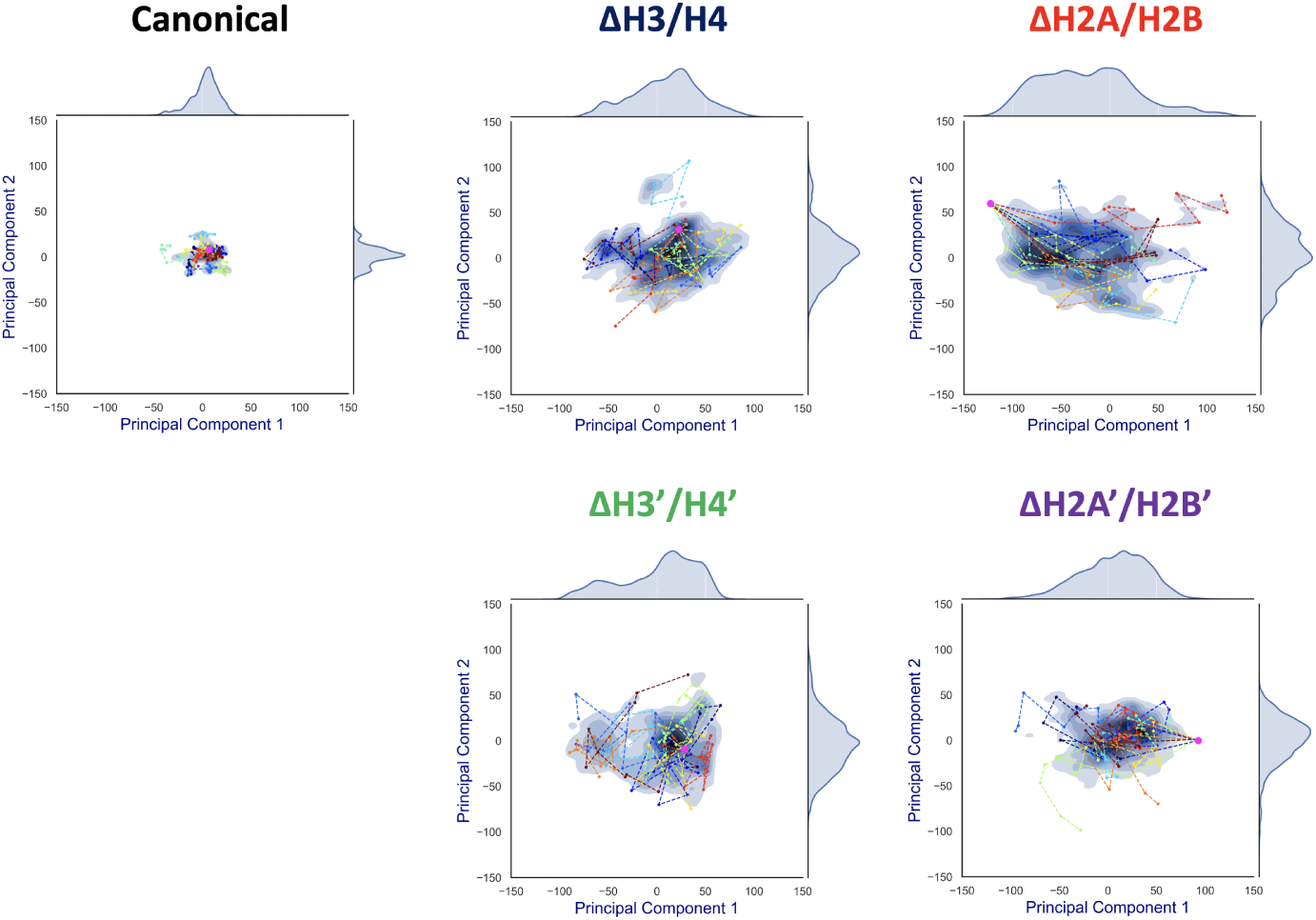
Free Energy Landscape of DNA Dynamics to 1st and 2nd PC Modes. The axes correspond to deviations (*S*(*t*, mode)) to the 1st and 2nd PC vector directions from the averaged structure (= *S*(*t*, 1) or *S*(*t*, 2)). The color (blue) shows the probability density of *S*. Each profile was fitted by kernel density estimation (KDE). Magenta dot corresponds to the initial conformation (*S*(0, 1), *S*(0, 2)), and each line (in different colors) shows a simulation trajectory (total 10; plotted at 10 ns intervals).

Contrasting results depending on the removed histone dimer were obtained in the PCA. In the case of nucleosomes lacking H2A/H2B (Δ H2A/H2B and Δ H2A’/H2B’), the initial structure (magenta point) was distant from the area with high probability density, and each simulation trajectory drifted from the initial structure (Fig. 5). This suggested that nucleosomes lacking H2A/H2B could not maintain their canonical histone-wrapped conformation, and deformed drastically. In contrast, in the case of H3/H4 lacking nucleosomes (Δ H3/H4 and Δ H3’/H4’), the initial coordinate (magenta point) locates around the high probability density area, and the trajectories were distributed around the initial coordinate (Fig. 5). While H3/H4 dissociation affects the nucleosome structure stability (Fig. 3), conformational changes often occur only around the initial structure.

### Interactions Between Histone Tails and DNA

We considered that the difference in the free energy profiles of DNA deformation was induced by physical restraints on DNA from histone proteins, and thus focused on physical interactions between DNA and histones. However, as shown in Fig. 3, structures of core histones changed only slightly. We also confirmed that the interaction between DNA and core histones did not show a clear difference. Hence, we evaluated contact frequency between the DNA and histone tails (Fig. S4). It is to be noted that the histone tails were not counted in the analysis of RMSD, RMSF, and PCA above (see Materials and Methods).

Comparing the contact matrices of the canonical nucleosome and others, no significant changes were observed in the majority of components. It is consistent with a previous study by MD simulations showing that histone tails were trapped by the adjacent DNA (28). However, in the nucleosomes lacking H3/H4 (Δ H3/H4 and Δ H3’/H4’), H2A C-terminal region adjacent to the dissociated H3/H4 showed enhanced contact with DNA (Blue rectangles in Fig. S4), which were not observed in the nucleosomes lacking H2A/H2B. Therefore, we concluded that the difference of free energy profiles (Fig. 5) were closely related to these histone tails.

Increase of contact frequency of nucleosomes lacking H3/H4 corresponds to the invasion of H2A C-terminal region to internal space originally occupied by the removed H3/H4 (Fig. 6). The H2A C-terminal interacts with the H3/H4 in the canonical nucleosome (1), and the contact frequency between the H2A C-terminal and DNA was increased by the invasion upon loss of H3/H4. These additional interactions restrict DNA deformation and breathing motion, and consequently the histone-wrapping DNA conformation was retained (Fig. 5). In contrast, in the nucleosomes lacking H2A/H2B, DNA neighboring the removed histone dimer shows drastic deformations (Fig. 6). The behavior of the H3/H4 tails is similar to previous study (29), and not altered by the loss of H2A/H2B.

**Figure 6:**
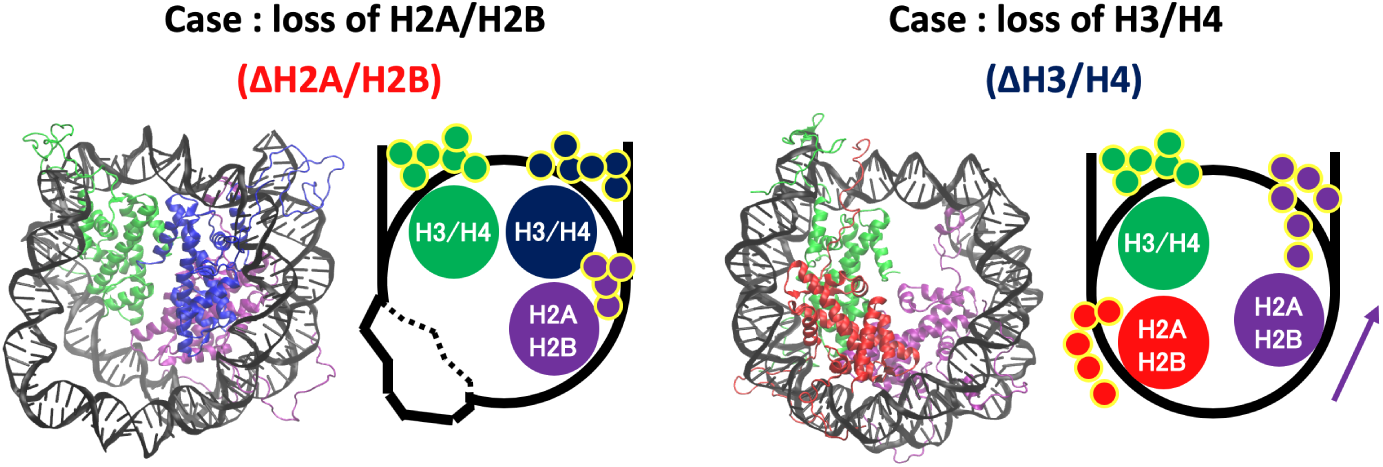
Histone Tail Dynamics in ΔH3/H4 and ΔH2A/H2B. Snapshots from simulation trajectories and schematics of Δ H3/H4 and Δ H2A/H2B are shown. Histone tails are represented by small circles in the schematics.

## Discussion

In the partially disassembled nucleosomes, the core histones showed no significant deformation. Indeed, RMSD of the hexasome (Δ H2A/H2B and Δ H2A’/H2B’), which was also experimentally observed, was very small (Fig. 3). We could not find any clear difference between the hexasome part in the overlapping di-nucleosome structure obtained by experiment (PDB ID: 5GSE) (15), and the nucleosome lacking H2A/H2B constructed from the canonical nucleosome structure (PDB ID: 1KX5). RMSD between them is 2.554 Å. Therefore, in partially disassembled nucleosomes, drastic structural deformation occurs only in the DNA, while histone structures are similar to the canonical crystal structure during attachment and detachment of histones. Note that unlike coarse-grained models depending on a reference structure, the force-field used in the all-atom simulations does not depend on a specific reference conformation.

On the other hand, DNA dynamics of nucleosomes lacking any histone dimer were statistically different depending on dissociated histones (Fig. 5). Individual histones did not show obvious structural deformation (Fig. 3). When an H2A/H2B was removed (Δ H2A/H2B and Δ H2A’/H2B’), N-terminal long tails of H3 were trapped at the dyad part (Fig. 6). In previous MD studies, it was shown that histone tails keep the DNA from being peeled off in the canonical nucleosome (28, 30–32). Particularly, the role of N-terminal tail of H3 in the stabilization was discussed. Our result is consistent with these studies. Due to the H3 tail trapping, free space induced by histone deletion was not filled, and DNA adjacent to the space was highly variable (Fig. 6). Similar DNA-histone interactions may exist after the H2A/H2B release induced by spontaneous breathing and partial unwrapping (27, 33), which may play a role in further nucleosome deformation and histone exchange. Although our simulations correspond only to the situation after H2A/H2B release and does not show its cause, the observed DNA fluctuations may help histone chaperones to intrude between DNA and histones. In contrast, when an H3/H4 was removed (Δ H3/H4 and Δ H3’/H4’), negligible deviation from the initial structure was observed (Fig. 5). DNA that originally interacted with the H3/H4 in turn interacted with the H2A C-terminal region adjacent to the H3/H4, which stabilized the DNA (Fig. 6). Due to the invasion and transient interaction of the H2A C-terminal, the nucleosome showed conformation relatively close to the canonical nucleosome. Single-molecule experiments using e.g. Förster resonance energy transfer (34) would be effective for verification of such changes in DNA dynamics.

From the results above, the following scenario for nucleosome remodeling was suggested. H2A/H2B dissociation induces only DNA deformation. Then, reinsertion of H2A/H2B into such a nucleosome would be relatively easy because of structural flexibility (Fig. S5). In contrast, H3/H4 dissociation induces additional interaction between DNA and H2A C-terminal tail (Fig. 6). This interaction prevents DNA deformation, and keeps the structure similar to the canonical one, which makes reinsertion difficult. For reversible attachment and detachment of histone proteins, H2A/H2B has a dynamic advantage. In fact, nucleosomal structure modulation by histone chaperones has been experimentally observed only in H2A/H2B dissociation. Such selectivity may arise from the physical property. It may also be relevant to localization of histone variants. CENP-A and H3.3, well-known histone variants of H3, localize in the centromere and telomeric region, respectively (35, 36). These variants show specific location in the whole genome and respective chromatin conformation. On the other hand, H2A.X, H2A.Z, and other H2A variants are distributed over the entire genome, and show various functions (17, 18, 37). This difference in distributions is also consistent with our study. As shown above, the exchange (i.e. attachment and detachment) of H3/H4 is infrequent. The substitution of H3 with another H3 variant is also difficult, resulting in the maintenance of the specific location in the whole genome. In contrast, substitution for H2A/H2B seems possible. Thus, their location and function do not show specificity. Hence, the observed nucleosome dynamics may also relate to the selectivity of histones and acquisition of biochemical functions.

## Conclusion

We analyzed dynamical behavior of partially disassembled nucleosomes using MD simulations. We observed that removal of an H2A/H2B dimer from the canonical nucleosome induces large fluctuations of nucleosomal DNA, while keeping the space for the histone dimer unoccupied. In contrast, when an H3/H4 was removed, the space for the dimer was invaded by the C-terminal tail of H2A, which mitigated the DNA fluctuations. The observed behavior may support frequent exchange of H2A/H2B, as well as specific molecular mechanisms such as histone chaperones.

## Author Contributions

Conceived and designed the research: TK AA YT. Performed the simulations: TK YT. Analyzed the data: TK AA YT. Wrote the paper: TK AA YT.

## Acknowledgments

The authors are grateful to K. Saikusa, R. Kawasaki, N. Sakamoto, and H. Nishimori for fruitful discussions. This work was supported by JSPS KAKENHI Grant Numbers JP17K05614 (to A.A.), JP16H01408 and JP18H04720, and by JSPS Bilateral Joint Research Project.

